# The *Vibrio vulnificus* stressosome is dispensable in nutrient-replete conditions

**DOI:** 10.1101/2022.01.27.477717

**Authors:** Laura Cutugno, Jennifer Mc Cafferty, Jan Pané-Farré, Conor O’Byrne, Aoife Boyd

## Abstract

The stressosome is a protein complex that has been demonstrated to sense environmental stresses and mediate the stress response in several Gram-positive bacteria, through the activation of the alternative sigma factor SigB. The i*n vivo* characterisation of this complex has never been performed in *Vibrio vulnificus* or any other bacteria that do not possess SigB. The elucidation of the role of the stressosome in *V. vulnificus* would provide elements to elaborate a functional model of the complex in a Gram-negative bacterium and identify the regulatory output in the absence of SigB. The stressosome locus is only found in 44% of *Vibrio vulnificus* isolates raising the question as to whether the role of stressosome is essential or modulatory in this bacterial species.

In this work, the expression of the stressosome genes was proven in nutrient-replete conditions and the co-transcription as one operonic unit of the stressosome locus and its putative downstream regulatory locus was demonstrated.

Moreover, the construction of a stressosome mutant lacking the four genes constituting the stressosome complex allowed us to examine the role of this complex *in vivo*. The initial established mutagenesis strategy relied on rifampicin-resistant *V. vulnificus* to select recombinant bacteria. Our data clearly showed that the influence of the Rif^R^ allele on stress and virulence characteristics overshadowed any effects of the stressosome. Therefore, we established an alternative mutagenesis strategy with a non-modified *V. vulnificus* parental strain and a DAP auxotrophic *E. coli* donor strain. Extensive phenotypic characterisation of the successfully-generated mutant in nutrient-replete conditions showed that the stressosome does not significantly contribute to the growth, of *V. vulnificus*. The stressosome did not modulate the response of *V. vulnificus* to the range of stresses tested – Ethanol, osmolarity, temperature, and salinity. Furthermore, the stressosome is dispensable for motility and exoenzyme production of *V. vulnificus*.

## INTRODUCTION

*Vibrio vulnificus* is a human foodborne pathogen that causes severe human infections, with a fatality rate higher than 50% in the United States (1). It populates coastal waters and is bioaccumulated by many bivalves, amongst which are oysters and other shellfish used for human consumption (2). Two different infection pathways have been described for this pathogen, gastrointestinal and wound infections (2–4). The first one occurs when the bacteria is ingested through the consumption of raw molluscs (1), the latter is caused by the contact of contaminated water with pre-existing wounds and it represents the most common infection route in the US (5–7). The fatality rate of the latter pathway is high but the severity of the infection generally depends on underlying health conditions in the patient (2). *V. vulnificus* is considered an opportunistic pathogen and severe infections have been observed in immunocompromised patients and in the presence of pathological conditions that increase the blood level of iron, such as liver disease (8, 9). Moreover, the distribution of this pathogen is dependent on environmental factors, such as water temperature and salinity this results in regionality of the cases of *V. vulnificus* infection (10, 11). A key step to understanding and predicting the distribution of the microorganism and the occurrence of the infection is the elucidation of the mechanisms that constitute the bacterial stress response.

The bacterial stress response is the set of physiological changes and molecular mechanisms that a bacterium puts in place to survive changes in the environment that would otherwise be lethal (12). The stress response is a fundamental step to persist in the environment and to guarantee a successful infection in the case of pathogenic bacteria (13, 14). It does require substantial changes in the physiology of the bacteria that are often in contrast with normal growth and propagation (15). For this reason, it is essential to regulate the stress response and limit such changes to the right time and environmental conditions. To do so, bacteria have developed complex regulation mechanisms, often characterised by a fast activation and equally important and efficient inactivation systems. To optimally coordinate the stress response with the outside world, bacteria have evolved signalling complexes that sense the stress and integrate the signal to modulate the cell response. Amongst these is the bacterial stressosome, a 1.8 MDa complex that has been discovered and extensively characterised in the Gram-positive *Bacillus subtilis* (16–18). In this organism and other Gram-positive bacteria, the stressosome has been found to activate the alternative sigma factor σ^B^ (SigB), following the sensing of environmental stresses (19). SigB in its active form binds to the RNA polymerase (RNAP) and promotes the expression of hundreds of genes involved in stress response (20).

Interestingly, the genetic locus encoding for the stressosome proteins has been found in several phyla and in bacteria that do not possess SigB, like *V. vulnificus* (21). In this organism, the stressosome locus contains an upstream and a downstream module. The first formed by the genes encoding for the three stressosome proteins VvrsbR, VvrsbS, VvrsbT (RsbR, RsbS and RsbT in *B. subtilis*) and the phosphatase VvrsbX (RsbX) and the latter encoding for a putative regulatory output of the stressosome, a two-component system (TCS) most likely involved in c-di-GMP hydrolysis (22). This locus has been identified in approximately 50% of the sequenced strains and its expression has been proved both in nature and laboratory conditions (23–25). Moreover, the heme-binding globin domain at the N-terminal of VvuR and the biochemical characterisation of these proteins from *V. brasiliensis* has shown a potential role in surviving or tolerating anaerobic conditions (26). This would be relevant both for the persistence of *V. vulnificus* in the environment and for survival during the infection process. For this reason, this work has been focused on characterising, for the first time *in vivo*, the role of the stressosome in this marine pathogen through the construction and phenotypical characterisation of a stressosome mutant, lacking the whole upstream module. This preliminary characterisation confirmed that the stressosome genes are expressed in LB + 2.5% NaCl (LBN), a rich media commonly used to culture this bacterium and focused on the effects of the stressosome on the growth and stress response of *V. vulnificus*, with particular attention to anaerobiosis and oxidative stress. Moreover, due to the potential role in regulating the levels of c-di-GMP, motility and other virulence traits were tested to elucidate the role of the stressosome in the virulence of *V. vulnificus*.

## MATERIAL AND METHODS

### Strains, plasmids and growth conditions

The clinical strain *Vibrio vulnificus* CMCP6 (27) was used for the construction of the stressosome mutant in this work (see below). The auxotrophic *E. coli* β2163 was used to perform bacterial conjugation (28). *Vibrio vulnificus* was cultured in lysogeny broth medium with an additional 0.4M NaCl (LBN) at 30°C or 37°C, as specified. LB medium supplemented with 0.3 mM 2,6-Diaminopimelic acid (DAP) was used to culture *E. coli* β2163 at 37°C. Overnight broth cultures were grown in 2 ml in 15 ml bacterial culture tubes, at 37 °C with agitation at 150 rpm. Specific growth conditions, different from the one indicated above, are described in the appropriate sections. All chemicals and reagents were supplied by Sigma, unless indicated otherwise.

### RNA extraction and one-step RT-PCR analysis

For RNA extraction, *V. vulnificus* wild-type was grown overnight in LBN at 30°C and 37°C. One volume of bacterial culture containing approximately 0.2 OD_600_ of cells was mixed with 2 volumes of RNAprotect (Qiagen) and processed according to the manufacturer’s instructions. The pellet was used for RNA extraction, within 24 h of treatment with RNAprotect. The pellet was resuspended in 200 μl of TE buffer (30 mM Tris-HCl, 1mM EDTA, pH 8.0) + 15 mg/ml lysozyme + 10 μl ready-to-use QIAGEN Proteinase K (20 mg/ml) and incubated 10 min at room temperature to achieve cell lysis. RNA extraction on the lysate was performed with RNeasy Mini Kit (Qiagen) following the manufacturer’s protocol. The total RNA was eluted in 30 μL RNase-free water and traces of DNA were removed using the TURBO DNA-free kit (Life Technologies), according to the manufacturer instruction. The concentration and quality of the extracted RNA were assessed with a NanoDrop spectrophotometer and only samples with a 260/280 ratio ≥ 1.9 were used for the RT-PCR protocol, and RNA integrity was verified on a 1.5% agarose gel. To assess the presence of the target mRNA a one-step RT-PCR protocol was performed, using the QIAGEN OneStep RT-PCR Kit, according to the manufacturer instruction. Briefly, the cycle consisted of two subsequent steps, one of reverse transcription (RT) at 50°C for 30 min and one traditional PCR cycle. The primers for the PCR reactions (Table 1) were designed using Primer-BLAST (29). An RT negative control, that skipped the step at 50°C, was included to confirm the absence of DNA contamination. A PCR negative control in which the template was substituted with PCR-grade water was included for each pair of primers. The gene *tuf* (elongation factor Tu) was used as endogenous control. PCR products were visualised on a 1.2% agarose gel.

**Table 1.**
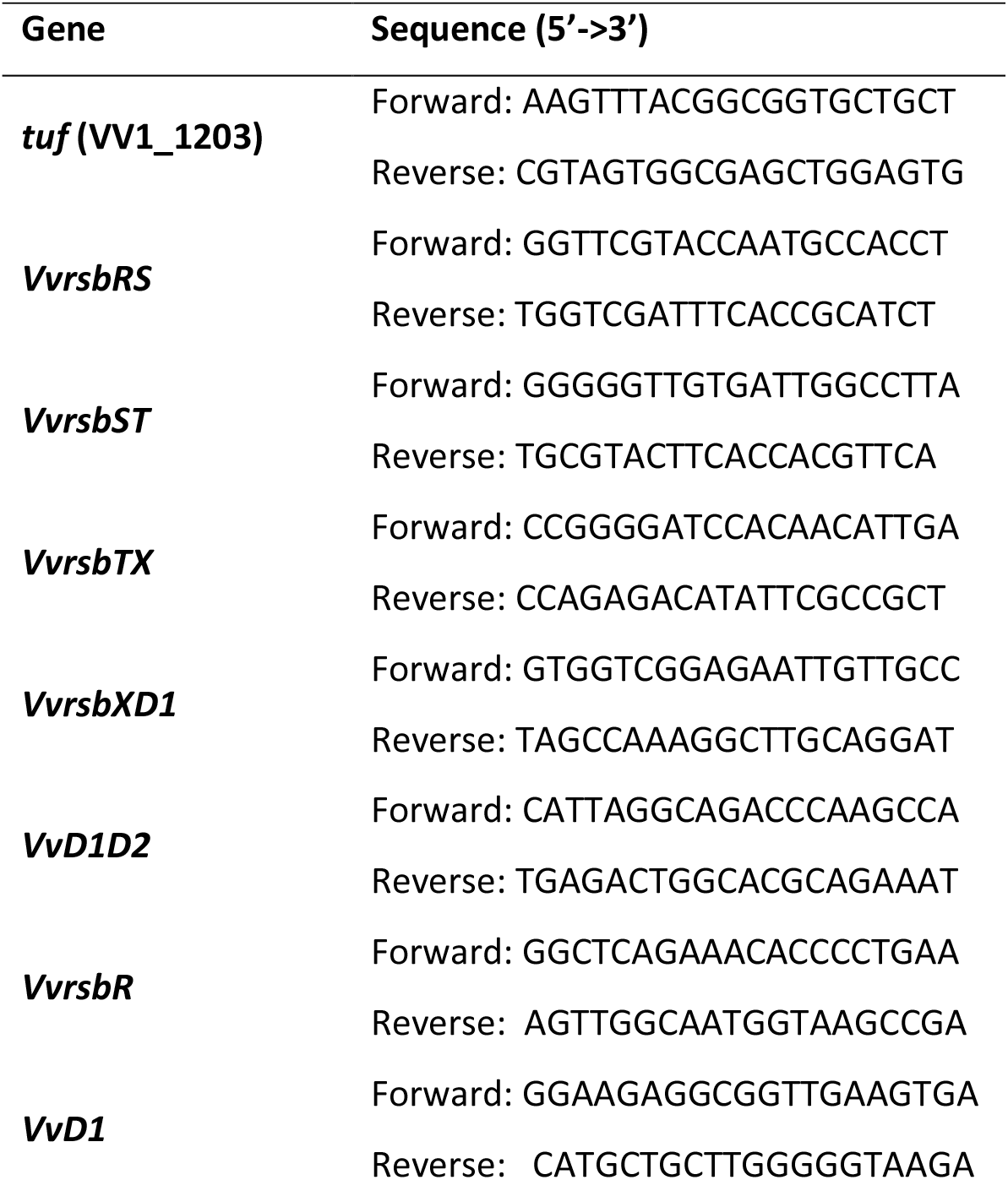
Primers for co-transcription experiments of the stressosome locus genes in *V. vulnificus* CMCP6 wild-type

### Construction of the *V. vulnificus* ΔRSTX knock-out mutant in a rifampicin-resistant *V. vulnificus* background

The construction of a knock-out mutant lacking the upstream module of the stressosome locus consisting of the *VvrsbR, VvrsbS, VvrsbT* and *VvrsbX* genes (*V. vulnificus* ΔRSTX), was first achieved using a classical conjugation protocol with a spontaneous rifampicin-resistant strain of *V. vulnificus* (Rif^R^) and *E.coli* SM10λpir. The use of a rifampicin-resistant *V. vulnificus* has been previously used (30, 31) and it is necessary to counter-select for the donor strain after conjugation. The knockout cassette (Eurofins) contained a deletion from nucleotide 4 of *VvrsbR* to nucleotide 577 of *VvrsbX*. The cassette was digested with *Xba*I and *Zra*I and cloned into the pDS132 suicide vector (32), between the *sacB* promoter and the *lac* operator, resulting in the knockout plasmids pDS_ΔRSTX. This was transformed by electroporation into *E. coli* SM10λpir and transformant colonies selected on LB + 25 μg/mL Chloramphenicol. Bi-parental conjugation was performed using the *E. coli* strain containing the pDS_ΔRSTX plasmid as a donor strain and the Rif^R^ *V. vulnificus* as parental strain. *V. vulnificus* CMCP6 and *E. coli* SM10λpir pDS_ΔRSTX were grown overnight at 37°C in LBN and LB + 25 μg/mL Chloramphenicol, respectively. Cultures were washed in LB broth, the two strains were mixed at a 1:1 ratio (v/v), spotted onto LB agar plates and incubated for 5h at 37°C. *V. vulnificus* transconjugants carrying the knock-out plasmid integrated into the chromosome by homologous recombination were selected on LBN + 5 μg/mL Chloramphenicol + 50 μg/mL Rifampicin. To promote the excision of the plasmid from the chromosome of *V. vulnificus*, the second event of homologous recombination was favoured by culturing the first recombinant cells in LBN broth without selective antibiotic. Second recombinant cells were selected on LBN + 5% sucrose (w/v) because the *sacB* gene present in the suicide plasmid does not allow growth in the presence of sucrose and growth on this media is a sign of a successful excision of the plasmid from the bacterial genome. This event can equally generate wild-type cells or deletion mutants. For this reason, mutants were selected by PCR screening of colonies grown on LBN + 5% sucrose using Taq polymerase (Bioline) and the primers RSTX_For (5’-GTCACGGGTTGATTGATTCGCAT-3’) and RSTX_Rev (5’-CTCACCGAGACGTAACATATGAATGT-3’). PCR products of different sizes for the mutant and the wild-type were visualised through agarose gel electrophoresis.

### Construction of the *V. vulnificus* ΔRSTX knock-out mutant in a wild-type *V. vulnificus* background

To avoid the use of a rifampicin-resistant strain of *V. vulnificus* and pleiotropic effects previously reported associated with such mutations (33) and observed also in this work to interfere with the phenotypic characterisation of the mutant, the construction of the stressosome knock-out mutant was performed in a wild-type background. To do so, we used for the first time a conjugation protocol with *V. vulnificus* and a DAP (diaminopimelic acid)-auxotrophic strain of *E. coli*. The same knock-out cassette (Eurofins) and plasmid described in the previous section were used and transformed by electroporation into *E. coli* β2163 and transformant colonies selected on LB + 25 μg/mL Chloramphenicol + 0.3 mM 2,6-DAP agar plates. Bi-parental conjugation was performed as previously described, with the only difference that 0.3 mM 2,6-DAP were supplemented to all the growth media to allow growth of the *E. coli* strain. Conjugation time was extended from 5h to 7h and *V. vulnificus* transconjugants were selected on LBN + 5 μg/mL Chloramphenicol. In this last medium, the absence of additional DAP allowed the counter-selection of the donor strain. Second recombination and selection of the mutant were performed as described in the previous section.

### Whole Genome Sequencing (WGS) analysis

One putative mutant was selected for each mutagenesis protocol and Whole Genome Sequencing (WGS) analysis was performed. In addition, the *V. vulnificus* CMCP6 and the Rif^R^ strain were also sequenced. Genomic DNA was extracted using the Wizard Genomic DNA Purification Kit (Promega), according to the manufacturer’s protocol. Genomic DNA was sequenced by MicrobesNG (Birmingham, UK) using Illumina technology. Average read lengths were between 168 and 645 nucleotides for each sample and average fold coverage was between 52 and 1162. BreSeq was used to call base substitution mutations Read Alignment evidence using Consensus mode, with a mutation E-value cut-off of 10 and frequency cut-off of 0.8 (80 %).

### Growth characterisation in LBN

Growth was assessed at 30°C in LBN, in the presence and reduced oxygen condition and at 42°C. Overnight cultures were grown in 2 mL LBN at 37°C and diluted in LBN at an initial OD of 0.01. 200 μL of cell suspension were incubated in a microtiter plate and incubated at the appropriate temperature. To achieve oxygen-depleted growth conditions a higher volume of cell suspension (approximately 300 μL) was used to completely fill the microtiter well and the plate was sealed with a sterile adhesive plastic film before incubation at the required temperature. To test growth at 42°C, the plate was set up as previously described and incubated first at 37°C for one hour and then at 42°C. At least three biological replicates were used for each strain and each of them was assessed in two technical replicates. The plates were statically incubated in a Sunrise microtiter plate reader at the specified temperature and the OD_595_ was measured every 30 min for 24h.

### Stress survival assays

In order to test stress survival in LBN, the survival of both *V. vulnificus* wild-type and the ΔRSTX mutant was assessed in LBN in the presence of different stresses. Overnights cultures were grown at 37°C in 2 mL LBN for no longer than 18h. The cultures were then diluted to OD 0.1 in 2 mL of the appropriate stress media and incubated at 30°C (except the samples tested for temperature stress). Stresses (and media compositions) were: ethanol stress (LBN + 10% ethanol (v/v)), oxidative stress (LBN + 2mM H_2_O_2_), acid stress (LBN pre-acidified to pH 4 with HCl) and temperature stress (45°C in a temperature-controlled block). 100 μL samples were withdrawn at different time points and used for serial dilution and plate counting. At least two biological replicates were tested for each strain and stress condition, each one assessed in two technical replicates for the plate counting.

### Stress tolerance assays

To test the ability of *V. vulnificus* wild-type and the ΔRSTX mutant to tolerate stress and to grow in non-lethal stress conditions, we tested the growth of the two strains on LBN agar supplemented with several stressors. Overnights cultures were grown at 37°C in 2 mL LBN for no longer than 18h and then diluted to OD 1 in LBN broth. Cells suspensions were then tenfold serial diluted up to 10^−7^ and 3 μL of each dilution spotted on the appropriate LBN agar plate and incubated at 30°C. Stresses (and media compositions) were: oxidative stress (LBN agar + 0.5 mM H_2_O_2_), osmotic stress (LB agar + 0.8 M NaCl). To test growth ability in complete anaerobic conditions, the plates were incubated in a 2.5L anaerobic jar in the presence of an Oxoid AnaeroGen 2.5L Sachet that generates an atmosphere with less than 1% oxygen, according to the manufacturer. Pictures of the plates were taken after 48h growth.

### Motility assay

In order to evaluate the effects of the stressosome mutation on the motility of *V. vulnificus*, the two strains were tested for swimming on rich motility agar plates (10 g tryptone, 20 g NaCl and 3.35 g agar per litre). Overnights cultures were grown overnight in 2 mL LBN broth at 37°C. A sterile metal wire was then immersed in the cell suspension and used to pierce the motility plate. Plates were incubated at 30°C or 37°C and motility zone measured after 16h. At least three biological strains were tested for each strain.

### Exoenzyme production

The wild-type and the ΔRSTX mutant strains were tested for the production of two exoenzymes, haemolysin and protease. The strains were grown overnight in 2 mL LBN broth at 37°C. The cell suspension was then inoculated on LBN agar plates containing 5 % (v/v) defibrinated sheep blood (ThermoFisher Scientific) for haemolysin assay and 1 % (w/v) skim milk for protease assay. The plates were incubated at 37°C and pictures were taken after 48 h and 24 h, for haemolysin and protease test respectively.

### Cross-protection assay

Cross-protection experiments were performed as previously described (34) to test the effects of nutrient downshift on temperature survival in *V. vulnificus* wild-type and ΔRSTX. Briefly, the strains were grown overnight in 2 mL LBN at 37°C. Cultures were then diluted in 2 mL of fresh LBN to OD_600_ 0.05 and grown to mid-log phase (OD_600_ 0.4-0.6). Cell cultures were then diluted 1:100 in CDM (9.94 mM Na_2_HPO_4_, 10.03 mM KH_2_PO_4_, 0.81 mM MgSO_4_·7H_2_O, 9.35 mM NH_4_Cl, 856 mM NaCl, 0.75 μM FeCl_3_) in the absence of carbon source. Cell suspensions were incubated at 37°C and 1 mL withdrawn at different time points (0h, 0.5h, 1h, 2h, 4h, 24h) to test survival to high temperature. Plate counting was performed, at each time point, before and after 1h of incubation at 45°C. Mid-log phase cells diluted in LBN rather than CDM were used as control, to confirm temperature sensitivity before nutritional downshift.

## RESULTS

### The stressosome locus is expressed in rich media with co-transcription of all modular genes

Previous reports have shown that *VvrsbR, VvrsbS, VvrsbT* and the downstream TCS (two-component system) are expressed in artificial seawater (ASW) (24) but little information is available on the expression and possible role of the stressosome in *V. vulnificus* during growth in rich media (35). Moreover, although the locus is predicted to be an operon, due to close proximity, sometimes overlapping, of the genes to one another, no proof of co-transcription has been provided to date. To address these two questions, the presence of single transcripts and co-transcripts was investigated in cells growing in LBN to stationary phase, through RNA extraction and non-quantitative Reverse Transcriptase PCR (RT-PCR). First, the presence of the transcripts of the *VvrsbR* and *VvrsbD1* genes was confirmed in cells growing in LBN, demonstrating that the locus is expressed in nutrient-replete conditions (Figure S1). Moreover, the detection of RT-PCR products with the use of primers in the proximity of intergenic regions at the 3’ and 5’ ends of adjoining genes (Figure 1A) indicated the presence of co-transcripts between each pair of genes within each module and between *VvrsbX* and *VvD1* (Figure 1B), showing that the upstream and downstream module genes are co-transcribed. The expression of the stressosome and its downstream module genes in LBN suggested a role of this complex in rich media, while their proximity on the chromosome and the suggested presence of a common mRNA support the idea of the two modules being functionally related to each other.

**Figure 1.**
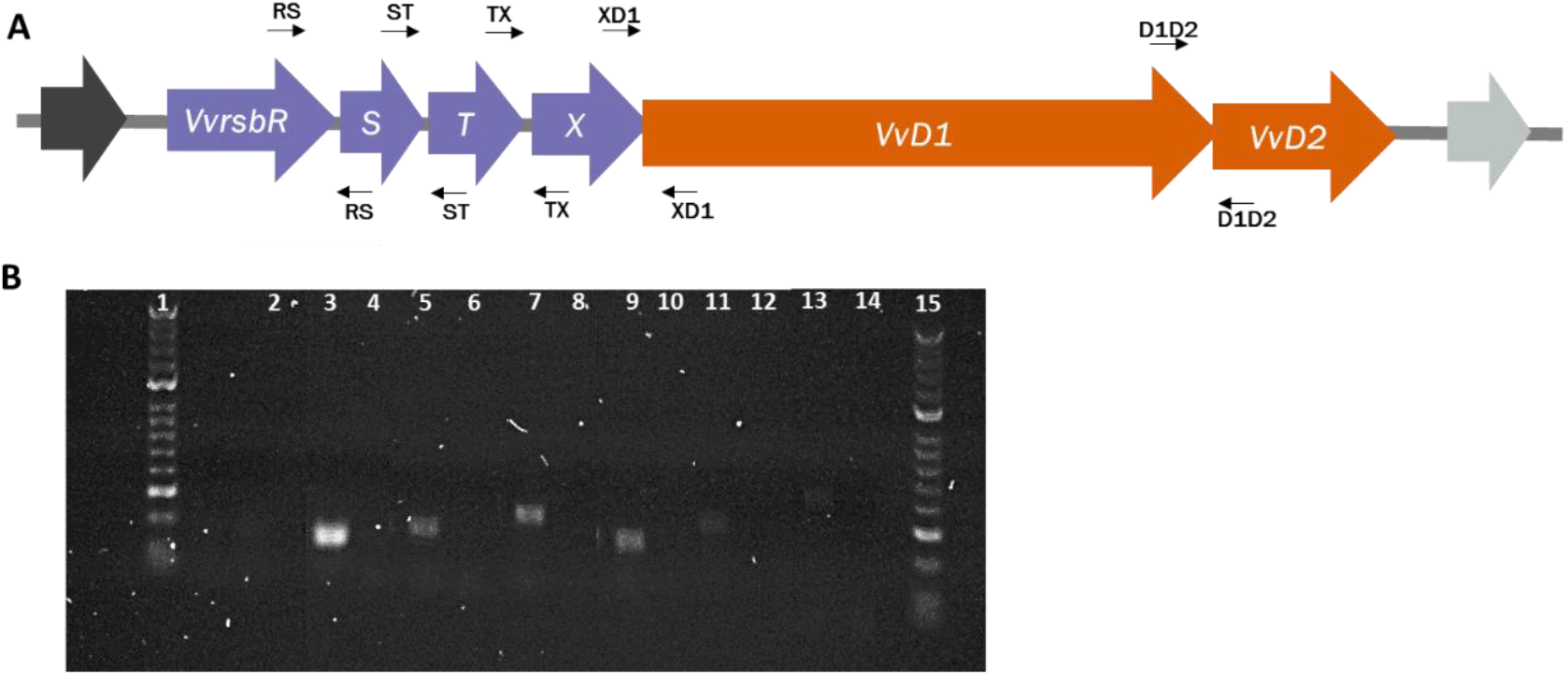
Co-transcription of stressosome locus genes in *Vibrio vulnificus*. A) The genetic stressosome locus in *V. vulnificus* CMCP6 wild-type, formed by the upstream module (*VvrsbR, VvrsbS, VvrsbT and VvrsbX*) in purple and the downstream module (*VvD1* and *VvD2*), in orange. The primers used for the co-transcription analysis are indicated at the top (forward primers) and at the bottom (reverse primers) of each gene. B) Co-transcription analysis of the stressosome locus genes. Electrophoresis analysis of the products of the RT-PCR performed on stationary phase cells, growing in LBN at 37°C. 1: HyperLadder 50bp (Bioline); 2: *tuf* RT negative control; 3: *tuf* gene; 4: *tuf* PCR negative control; 5: *VvrsbR-VvrsbS* co-transcript; 6: *VvrsbR-VvrsbS* PCR negative control; 7: *VvrsbS-VvrsbT* co-transcript; 8: *VvrsbS-VvrsbT* PCR negative control; 9: *VvrsbT-VvrsbX* co-transcript; 10: *VvrsbT-VvrsbX* PCR negative control; 11: *VvrsbX-VvD1* co-transcript; 12: *VvrsbX-VvD1* PCR negative control; 13: *VvD1-VvD2* co-transcript; 14: *VvD1-VvD2* PCR negative control; 15: HyperLadder 50bp (Bioline).

### Phenotypic characteristics of the stressosome mutant were overshadowed by the pleiotropic effects of the Rif^R^ allele

Phenotypic characterisation of the stressosome mutant constructed in the *V. vulnificus* rifampicin-resistant background was performed in order to identify the *in vivo* role of the stressosome in this human pathogen. The mutant was tested for growth, stress survival and tolerance, and for the main virulence characteristics and compared to the *V. vulnificus* CMCP6 wild-type strain. The use of the rifampicin-resistant parental strain as control, allowed us to identify possible effects of the rifampicin resistance and validate the use of the classical mutagenesis protocol when downstream applications include stress response and virulence characterisation of the mutants. In some cases, the mutant showed differences compared to the wild-type strain but all ascribable to pleiotropic effects in the rifampicin-resistant parental strain. In particular, reduced motility, ethanol survival and hyperosmotic stress tolerance were observed (Figure S2). The same effects were observed, to the same extent, in the rifampicin-resistant parental strain in line with previously published work (33). This is an indication that the construction of the stressosome mutant in a Rif^R^ strain might not allow the observation of differences caused by the *VvrsbRSTX* deletion, where these overlap with pleiotropic effects associated with rifampicin resistance. Based on this, a different mutagenesis protocol was successfully optimised during this work with the successful construction of the stressosome mutant in a wild-type background. This allowed a direct comparison with the wild-type and extensive characterisation of the role of the stressosome in LBN medium.

### Knockout mutation of the stressosome does not influence growth of *V. vulnificus* in LBN, at various oxygen concentrations and temperatures

In order to test the physiological role of the stressosome in *V. vulnificus*, a stressosome mutant lacking the upstream module (*V. vulnificus* ΔRSTX) was successfully constructed and used for phenotypic characterisation in rich media in comparison to the wild-type strain. Firstly, growth characterisation of the two strains was performed in LBN at 30°C and 37°C in aerobic conditions, to eliminate any possible influence of differential growth rate on the phenotypic characteristics analysed. The 2 strains grew with similar kinetics in these conditions (Fig S3)

Next, growth characterisation of the two strains was performed in LBN at 30°C and 37°C, in aerobic and O_2_-depleted conditions. *V. vulnificus* experiences variation in O_2_ levels both in the environment and in the human host (25). Moreover, the *V. brasiliensis* stressosome has been shown to bind O_2_ *in vitro* (26). For these reasons, growth curves in aerobic and reduced O_2_ conditions at 30°C (Figure 2A) were compared. Interestingly, the oxygen depletion does not cause any growth rate variation in the wild-type strain, but only a reduction in the overall biomass accumulation, indicating a good adaptation of *V. vulnificus* to hypoxia. Moreover, the growth profile of the mutant was comparable to the one of the wild-type strain, thus indicating no significant role of the stressosome in this adaptation to low oxygen levels.

**Figure 2.**
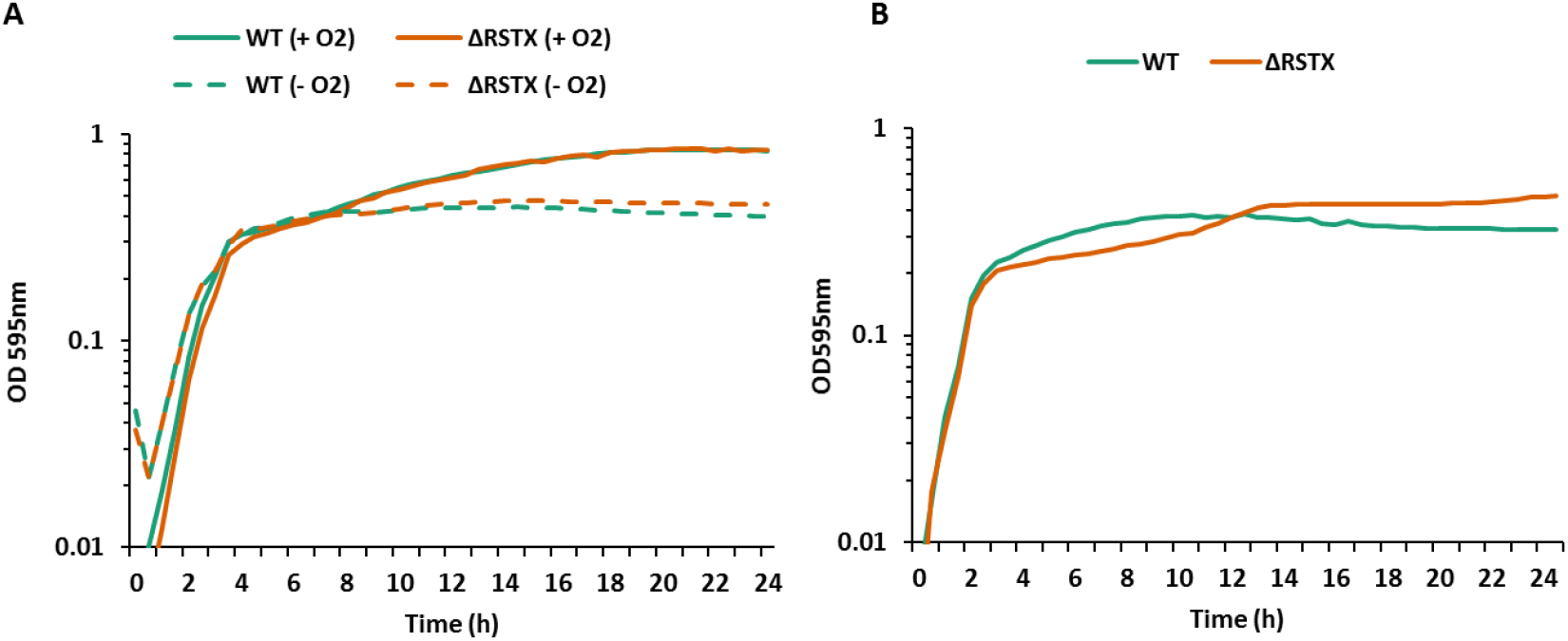
Growth characterisation of *V. vulnificus* CMP6 wild-type (green) and the ΔRSTX mutant (orange) in LBN. A) Growth curve in LBN at 30°C, in the presence of oxygen (continuous line) and in reduced oxygen conditions (dashed line). OD_595_ was measured every 30 min for 24 h. Each curve is the mean of three biological replicates. B) Growth curve in LBN at 42°C, after 1h growth at 37°C. OD_595_ was measured every 30 min for 24 h. Each curve is the mean of three biological replicates. Y-axis starts at the OD value of the inoculum (0.01).

During the infection of the human host, *V. vulnificus* often faces not only reduced oxygen levels but also increasing temperatures due to the occurrence of fever in the patient (36). For this reason, the ability of the two strains to grow at 42°C was tested after a brief time of growth at 37°C (Fig 2B). In this experimental setup, both the wild-type and the mutant strain showed slightly reduced growth when compared to the optimal growth temperature but no significant differences were observed between the two strains. These growth experiments demonstrate that the presence of the stressosome does not influence growth rates in LBN, nor is it required for the processes of adaptation to low oxygen levels or high temperatures in nutrient-replete conditions.

### The stressosome mutation does not alter the ability of *V. vulnificus* to survive lethal environmental stress in LBN

In Gram-positive bacteria, such as *Listeria monocytogenes* and *Bacillus subtilis*, the stressosome is part of the signalling hub that ultimately leads to the activation of the alternative sigma factor σ^B^ (22). This controls the transcription of hundreds of genes involved in the stress response and contributes to the survival of the bacteria in harsh environmental conditions (37–39). To study a possible role of the stressosome in the survival of *V. vulnificus* to lethal stresses, survival assays in the presence of a range of different stressors were performed using the wild-type *V. vulnificus* CMCP6 strain and the ΔRSTX strain (Figure 3). We tested several stresses, including 10% ethanol (Figure 3A), 2 mM H_2_O_2_ (Figure 3B), pH 4 (Figure 3C) and high temperature (45°C) (Figure 3D). Ethanol was used to test whether the difference observed for the Rif^R^ stressosome mutant was due to the rifampicin resistance and not to the *VvrsbRSTX* mutation. Oxidative, acid and temperature stresses were chosen because they are relevant to the life cycle of *V. vulnificus* and in particular to survive in the human host. Moreover, response mechanisms to these stresses have been only partially elucidated in *V. vulnificus* (40–42). The experiments performed showed that the two strains equally survived the tested stresses. In particular, no difference at all was observed in the presence of 10% ethanol and at pH 4, confirming that the use of the Rif^R^ parental strain was the cause of the previously observed difference. A faster death was occasionally observed in the presence of H_2_O_2_ (Figure 3B) and after a shift to 45°C (Figure 3.D). This might have a biological meaning and indicate a potential modulatory involvement of the stressosome or simply be due to overall higher variability in these stress conditions.

**Figure 3.**
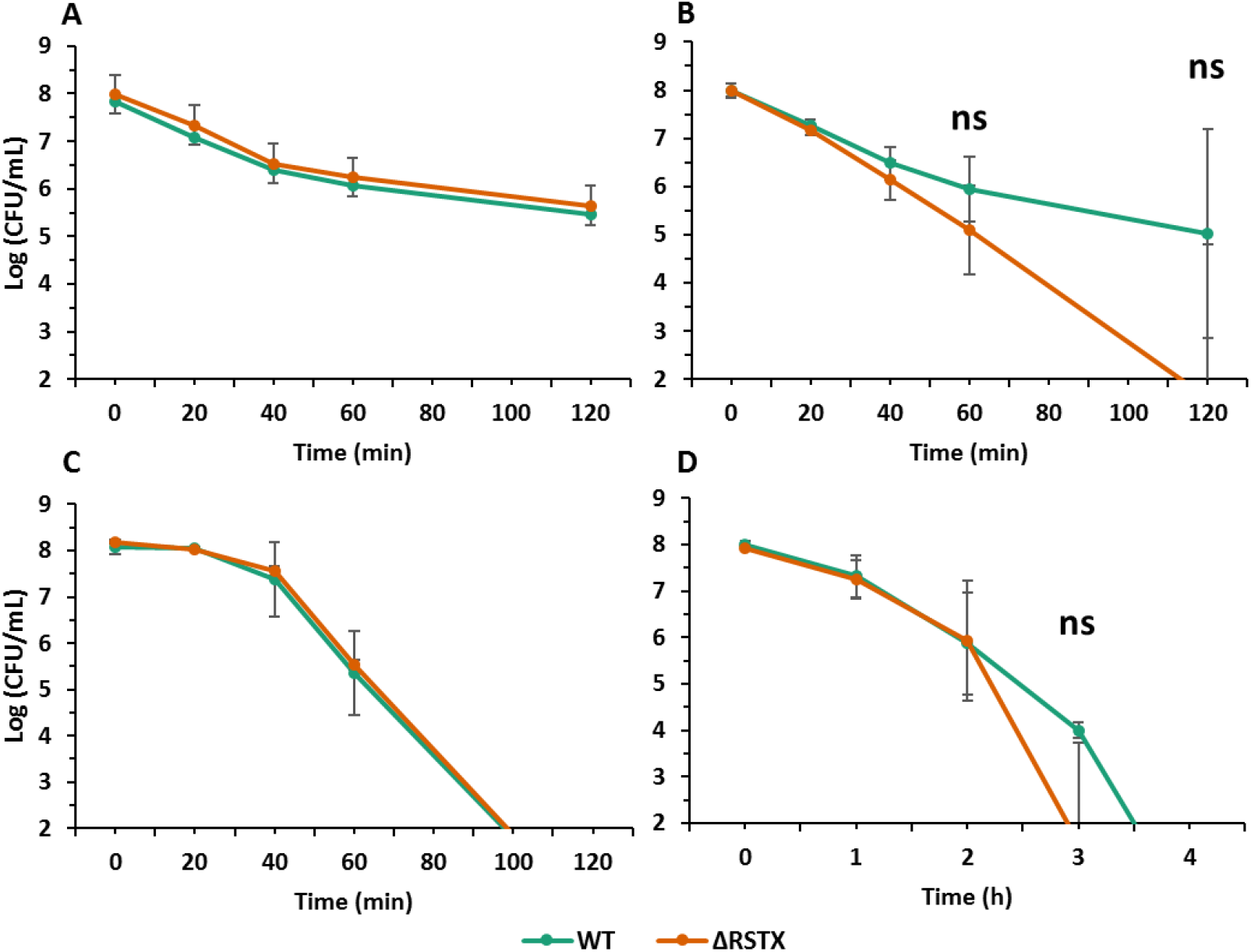
Stress survival of *V. vulnificus* CMP6 wild-type (green) and the ΔRSTX mutant (orange) in LBN. A) Survival assay in LBN + 10% Ethanol (v/v). The strains were incubated at 30°C and survival was assessed, through plate counting, at five different time points (0 min, 20 min, 40 min, 60 min and 120 min). B) Survival assay in LBN + 2 mM H_2_O_2_. The strains were incubated at 30°C and survival was assessed, through plate counting, at five different time points (0 min, 20 min, 40 min, 60 min and 120 min). C) Survival assay in LBN at pH 4. The strains were incubated at 30°C and survival was assessed, through plate counting, at five different time points (0 min, 20 min, 40 min, 60 min and 120 min). D) Survival assay in LBN at 45°C. The strains were incubated at 45°C and survival was assessed, through plate counting, at five different time points (0h, 1h, 2h, 3h and 4h). For each time point, survival was plotted as Log10(CFU/ml). Three biological replicates for each strain were tested and the reported values are the mean of the three replicates. Student’s t-test was performed comparing mutant strains to WT and P values are shown (ns > 0.05). Y-axis starts at Log (CFU/mL) = 2, in proximity of the detection limit value (2.4).

Higher temperatures (48°C and 50°C) and hyperosmotic stress (LBN + 1.2 M NaCl) were also tested but these were respectively too harsh or too mild conditions to appreciate differences between the strains (data not shown). In fact, the temperatures tested caused the complete death of both strains in less than 20 min thus making it challenging to perform a time-series experiment. The presence of an additional 1.2 M NaCl, on the contrary, did not cause bacterial death. These results indicate that, in our experimental conditions, the stressosome does not contribute to the survival of *V. vulnificus* to environmental stresses and its deletion does not affect the ability of this human pathogen to survive several stresses that can be encountered both in the environment and in the human host.

Concerning stress survival, several cases of cross-protection mechanisms have been described in *Vibrio* spp. (43) and in *V. vulnificus* specifically (34, 44), where pre-exposure to sub-lethal stresses (such as nutrient downshift) causes a general stress adaptation response that results in a significant increase in survival to subsequent exposure to lethal conditions. In this work, the role of the stressosome in the general stress adaptation response was assessed by determining survival to high temperature following nutrient downshift, well-described in *V. vulnificus* (34). We confirmed that the shift from a rich to a chemically defined media, lacking a carbon source, rapidly caused a significant transient resistance to lethal heat stress (1h at 45°C) but no differences were observed between the wild-type and the stressosome mutant (Figure 4). The cross-protective adaptive response occurred within 5 minutes following nutrient downshift and persisted for 2 h, after which time the adaptive response was resolved and normal responses returned. These experiments suggest that the stressosome does not regulate stress survival, neither directly nor indirectly through cross-protection adaptive mechanisms.

**Figure 4.**
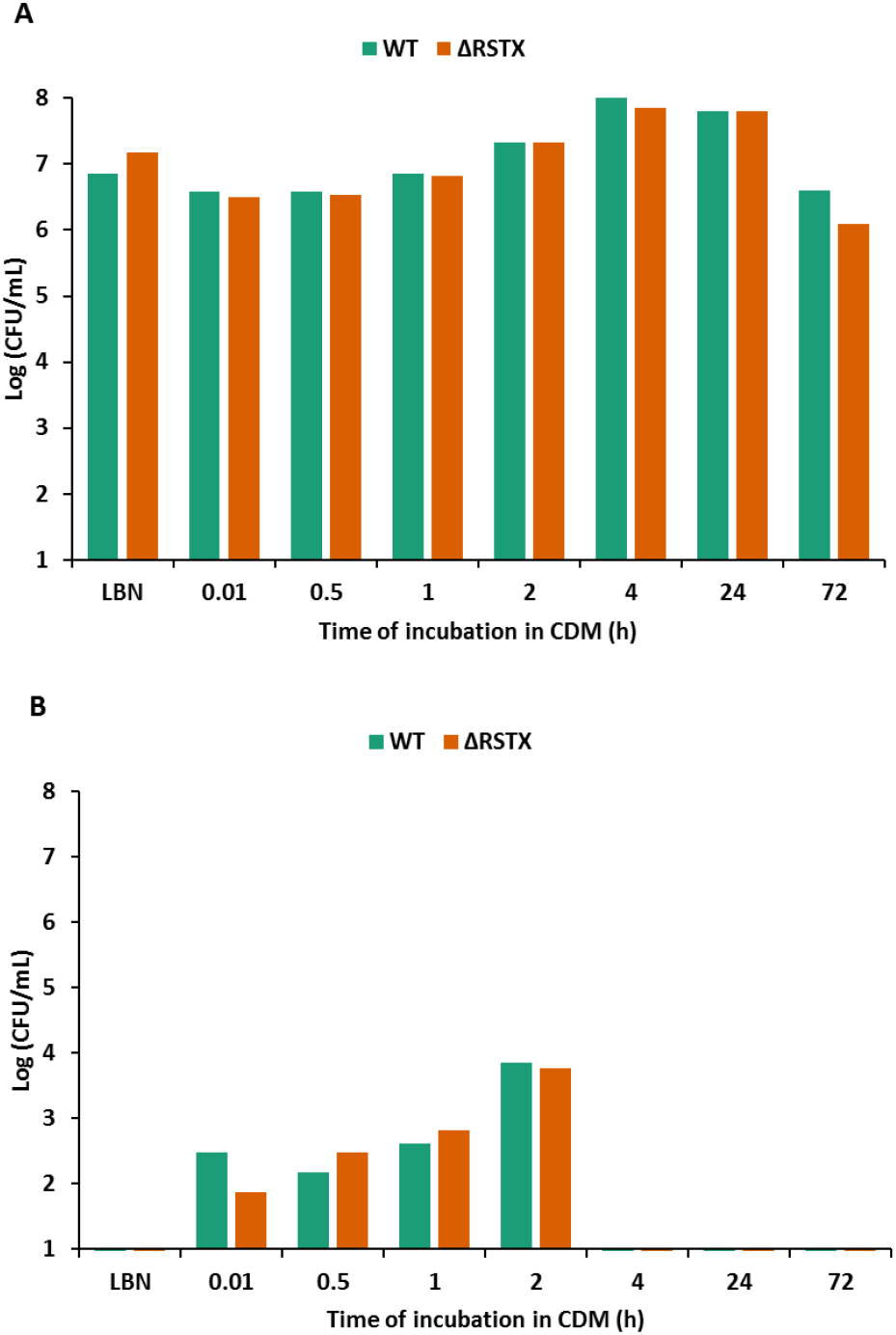
The cross-protection mechanism in *V. vulnificus* CMCP6 wild-type (green) and the ΔRSTX mutant (orange). A) Cell viability in LBN and CDM + 0.75 μM FeCl_3_ in the absence of glucose. The strains were incubated at 37°C and survival was assessed, through plate counting, before the shift from LBN to CDM and at six different time points (0 h, 0.5 h, 1h, 2h, 4h and 24h). A) Survival in LBN at 45°C for 1h. The strains were used for survival assay before the shift from LBN to CDM and at six different time points (0 h, 0.5 h, 1h, 2h, 4h and 24h). Survival to high temperature was assessed, through plate counting, after 1h at 45°C.

### The stressosome does not modulate stress tolerance in *V. vulnificus* in nutrient replete media

Bacterial stress response refers not only to survival to extreme lethal stresses but also to stress tolerance mechanisms that allow the microorganisms to grow in the presence of milder stresses. This is essential to allow a bacterium or a bacterial community to reproduce and grow in the environment, which must be considered as a dynamic system in which small changes happen continuously due to natural fluctuations, human intervention and the presence of other organisms and microorganisms (45). To test the ability of our strains to tolerate, adapt and grow in the presence of mild stresses, we analysed their growth on LBN agar in the presence of different stressors. Amongst these, we tested 0.5 mM H_2_O_2_, anaerobiosis and hyperosmotic stress (LB + 0.8 M NaCl) (Figure 5). Various degrees of growth effects were observed in the presence of these stresses for the wild-type, with the 0.5 mM H_2_O_2_ being the one causing the strongest inhibitory effect, with a reduction of five log in CFU numbers when compared to the LBN plate. Also the presence of extra salt caused a significant reduction in the growth ability of *V. vulnificus*, while the anaerobiosis condition only caused an effect on the amount of biomass present (i.e. smaller colonies indicative of a reduced replication rate) but no changes in the ability to initiate growth (i.e. equal number of colonies). The stressosome mutant strain suffered growth reductions comparable to those of the wild-type in all tested conditions. This confirms that the stressosome is not involved in the stress response of *V. vulnificus* in nutrient-replete conditions.

**Figure 5.**
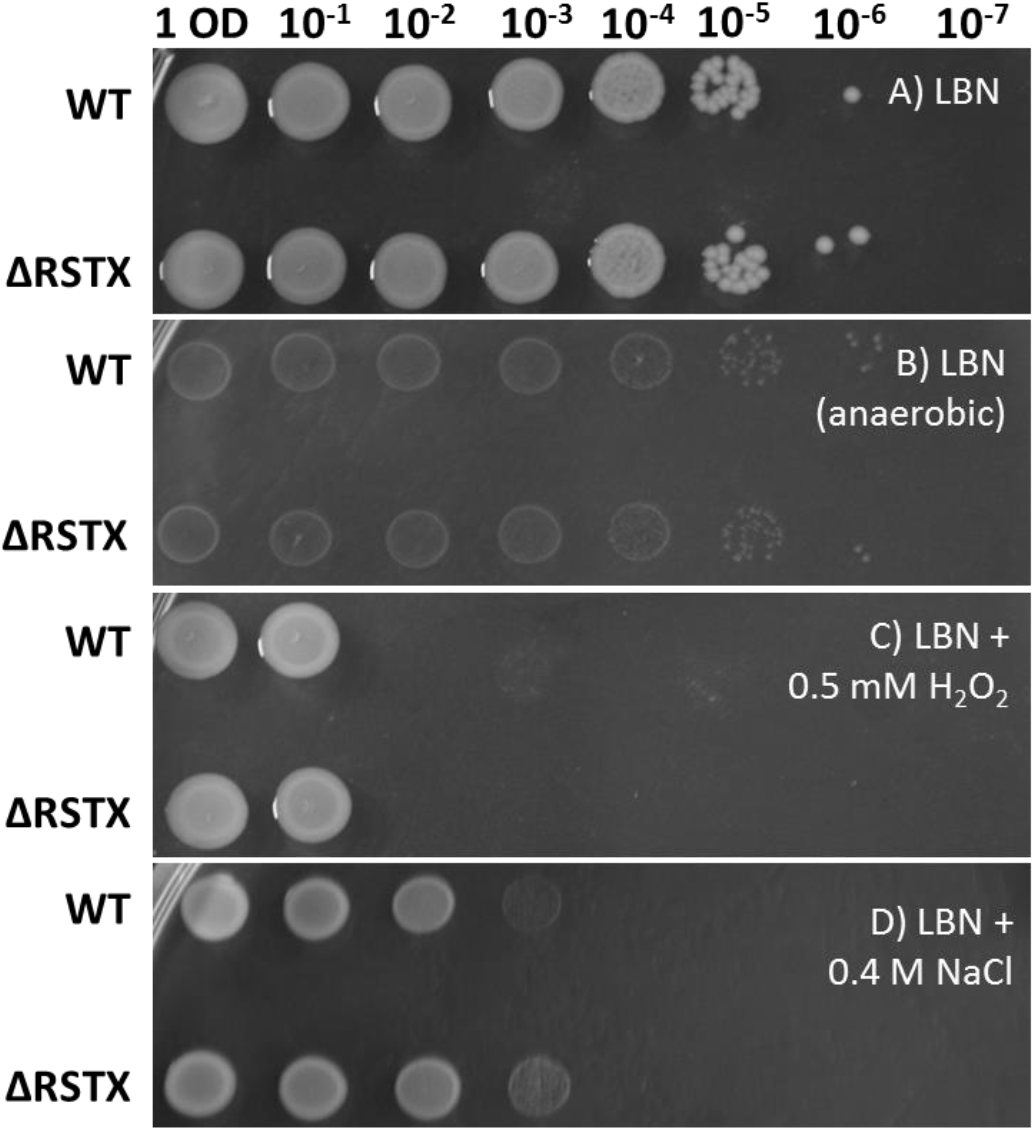
Growth assessment in stress conditions of *V. vulnificus* CMCP6 wild-type and the ΔRSTX mutant in LBN. The strains were first diluted to OD_600_ = 1 and then 10-fold serial dilutions were performed up to 10^−7^ and each dilution was spotted on agar plates and incubated at 30°C. A) LBN agar (LB agar + 0.4M NaCl) after 24 h incubation in atmospheric condition. B) LBN agar after 24 h incubation in oxygen-depleted condition. C) LBN agar + 0.5 mM H_2_O_2_ after 24 h incubation in atmospheric condition. D) LB agar + 0.8 M NaCl after 24 h incubation in atmospheric condition. The pictures shown are representative of at least two biological replicates.

### The stressosome does not affect motility and exoenzyme production

The previous experiments indicate the lack of a significant role of the stressosome in regulating growth and stress response in *V. vulnificus* in nutrient-replete media, in terms of stress survival, stress adaptation and stress tolerance. This leaves an open question on the physiological role of this complex in *V. vulnificus* that can explain the presence of the genetic locus in a large proportion of strains of this species and active transcription of the locus in LBN. To address this question, we investigated another fundamental aspect of the life cycle of a human pathogen: virulence characteristics. Virulence includes a wide range of physiological processes from motility to biofilm formation, through exoenzyme and toxin production, and it is intrinsically connected to both growth and stress response (46, 47).

In this work, we assessed the ability of both strains to swim on tryptone motility agar plates and their ability to produce haemolysin and protease exoenzymes (Figure 6). The motility was tested at both 30°C and 37°C (Figure 6A) and the results show a clear effect of the temperature but no effect of the stressosome mutation on the swimming of *V. vulnificus*. Haemolysin (Figure 6B) and protease (Figure 6C) production were tested at 37°C on LBN supplemented with sheep blood and skim milk respectively. Both *V. vulnificus* wild-type and ΔRSTX showed degradation of the substrates, indicating that they are actively producing and secreting both haemolysin and protease. Although precise quantification is not possible using this method, no significant difference was observed between the two.

**Figure 6.**
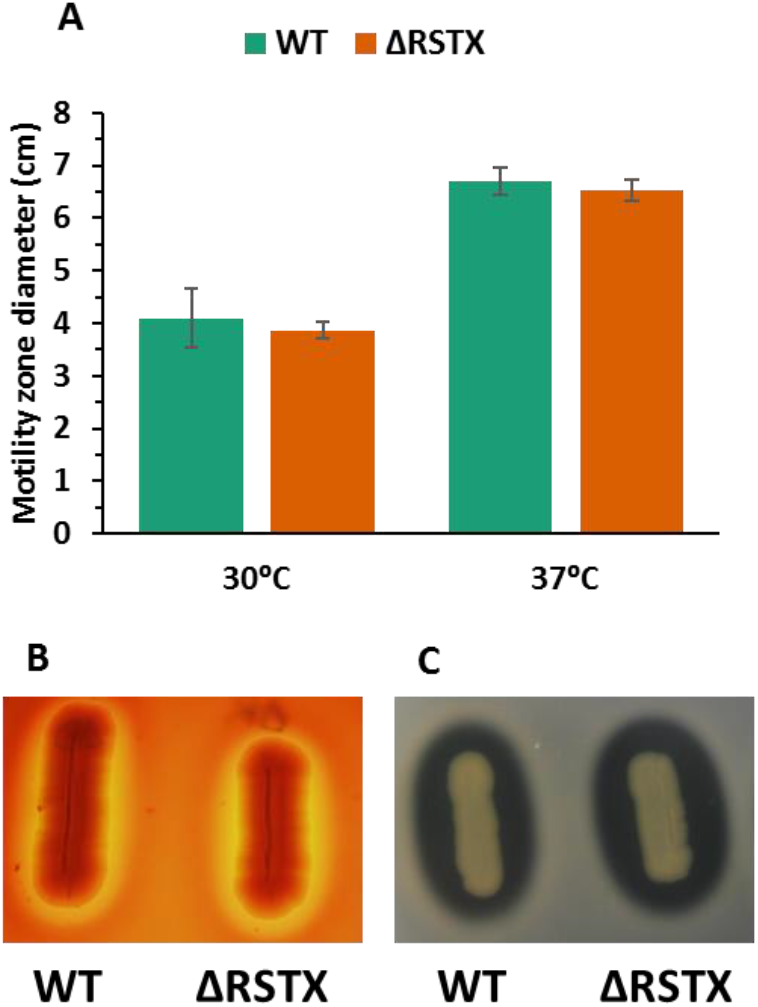
Motility and exoenzymes production characterisation of *V. vulnificus* CMCP6 wild-type (green) and the ΔRSTX mutant (orange) in rich media. A) Motility assay on tryptone motility plates. The strains were stabbed on motility agar plates and incubated at 30°C and 37°C. The motility zone was measured after 16h incubation. Three biological triplicates for each strain were tested and the reported values are the mean of the three replicates. B) Haemolysin production was tested on LBN agar plates + 5% sheep blood (v/v). Pictures were taken after 48h incubation at 37°C. C) Protease production was tested on LBN agar plates + 1% skimmed milk. Pictures were taken after 24h incubation at 37°C.

This set of experiments showed that the stressosome is not involved in motility and exoenzyme production in nutrient-replete conditions.

## DISCUSSION

The elucidation of the role of the stressosome in *V. vulnificus* could constitute an important piece of information towards the understanding of distribution and infection of a deadly and enigmatic human pathogen. Moreover, it would represent the first characterisation of the stressosome in a Gram-negative bacterium. This would lay the basis to elaborate a functional model of the stressosome different from the one observed in *B. subtilis* and other Gram-positive bacteria and to understand the evolution of this complex across different bacterial species (22).

In this work, we focused on the phenotypical characterisation of a stressosome mutant lacking the whole upstream module of the stressosome genetic locus. This includes the genes encoding the three proteins forming the stressosome (VvrsbR, VvrsbS and VvrsbT) and the phosphatase VvrsbX that has been shown to switch off the stressosome-mediated response and avoid the activation of SigB in the absence of stress in other microorganisms (48, 49). We decided to perform this extensive characterisation in the rich medium LBN. This is one of the most used media to study the physiology of *V. vulnificus* and it prevents the occurrence of nutritional stress, thus allowing us to focus exclusively on the environmental stress response. This is the route of SigB activation that is known to be mediated by the stressosome in *B. subtilis* (19).

We first confirmed the presence of the stressosome mRNA in wild-type cells grown to stationary phase in LBN, as the expression of these genes has been shown before only in ASW (24). We also investigated the possible co-transcription of the upstream and downstream modules and we confirmed that products of co-transcription were present. This, together with the co-localisation on chromosome two of *V. vulnificus*, supports the hypothesis that the two modules are functionally related and the downstream TCS could be the regulatory output of the stressosome in this bacterium.

Following this, construction of the knockout mutant using a classical mutagenesis protocol in a rifampicin-resistant parental background, led to the observation that pleiotropic effects of *rpoB* mutations causing rifampicin resistance can interfere with stress response and virulence characterisation of *V. vulnificus*. For this reason, we successfully optimised a new mutagenesis protocol that allowed construction of the mutant in a wild-type background, thanks to the use of an auxotrophic *E. coli* donor strain. The mutant constructed in the wild-type background was then utilised for an extensive phenotypical characterisation in comparison to the wild-type strain. This started with a growth characterisation in LBN. We confirmed no differences in the absence of stress, as expected, and focused on the effects of hypoxia and high temperature on growth. Differences between the two strains did not emerge and effects of hypoxia and high temperature were comparable between the two strains. Due to the predicted role of the stressosome in stress sensing and stress response, the next step of this phenotypical characterisation was the assessment of stress survival and tolerance of the stressosome-lacking mutant. Survival was assessed in the presence of several stresses, including acid, ethanol, oxidative and temperature stress, and once again different effects were observed on the wild-type but no differences emerged with the stressosome mutant. Stress tolerance, defined as the ability to grow in the presence of sublethal stresses, was also tested and no effects of the stressosome were observed. According to these results, the presence of the stressosome genes confers no advantage to the human pathogen *V. vulnificus* in growth, stress tolerance and stress survival in LBN. This is surprising considering the previously characterised role of the stressosome in Gram-positive bacteria (18, 22) and the presence of the locus transcript in these experimental conditions.

This lack of detectable function in mediating growth and survival in the presence of sublethal and lethal stresses led to the hypothesis that physiological roles of the stressosome could involve the colonisation and infection process. For this reason, motility and exoenzyme production of *V. vulnificus* wild-type and ΔRSTX mutant were tested. Again, the lack of the stressosome did not affect the ability of *V. vulnificus* to swim and produce enzymes relevant to the infection process, in our experimental condition.

These results could indicate a lack of function of the stressosome in *V. vulnificus* and explain why the stressosome locus is present only in 50% of the sequenced strains. On the other hand, it is possible that the experimental conditions used were not compatible with a stressosome activation, despite the presence of the locus transcripts. A basal level of stressosome expression may occur as a means for the bacterium to monitor and survey for the emergence of stress signal inducers. This could explain the detection of transcription of the stressosome genes in non-stressful conditions. This could be seen as been analogous to the role Toll-Like Receptors in eukaryotic cells acting as surveillance mechanisms for Pathogen-Associated and Damage-Associated Molecular Patterns that upon recognition and activation then initiate signalling cascades leading to inflammation and other cellular responses. Previous works have shown that the stressosome genes are expressed in ASW more than in human serum (24), thus indicating that the stressosome-mediated response could be naturally active in a nutrient-deficient environment rather than in rich media. This first *in vivo* extensive characterisation in a rich medium is the demonstration that the stressosome does not affect *V. vulnificus* physiology in a nutrient-replete environment. Based on this and previous transcriptional analysis, we suggest that a similar characterisation in a minimal, nutrient-depleted media should be performed to identify the role of the stressosome and identify the putative trigger and regulatory output of this complex in a Gram-negative human pathogen.

## ACKNOWLEDGEMENT

This research was jointly supported by a Marie Skłodowska-Curie ITN Fellowship (PATHSENSE, project number 721456) within the seventh European Community Framework Program and by the Irish Research Council (IRC), under grant number GOIPG/2020/1420. Additional support was provided by the Irish HEA (Higher Education Authority) as part of a COVID-19 costed extension programme.

Genome sequencing was provided by MicrobesNG (http://www.microbesng.uk) which is supported by the BBSRC (grant number BB/L024209/1).

We thank Prof Miguel A. Valvano (Queen’s University Belfast) for kindly providing the *E. coli* β2163 strain.

## CONFLICTS OF INTEREST

The authors declare that there are no conflicts of interest.

## Notes

### Competing Interest Statement

The authors have declared no competing interest.

