## Supplementary Figures S1, S2 and S3 for "The *Vibrio vulnificus* stressosome is dispensable in nutrient-replete conditions"

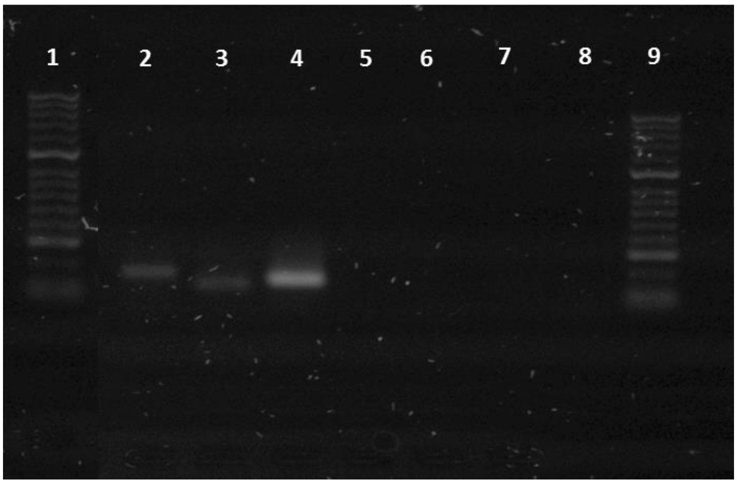


**Figure S1. Transcription of stressosome locus genes in stationary-growing cells of *Vibrio vulnificus* in LBN.** RT-PCR analysis of the *VvrsbR* and *VvD1* genes. Electrophoresis analysis of the products of the RT-PCR performed on stationary phase cells, growing in LBN at $37^{o}$C. 1: HyperLadder 50bp (Bioline); 2: *VvrsbR* gene transcript; 3: *VvD1* gene transcript; 4: *tuf* gene transcript; 5: *VvrsbR* PCR negative control; 6: *VvD1* PCR negative control; 7: *tuf* PCR negative control; 8: *tuf* RT negative control; 9: HyperLadder 50bp (Bioline).

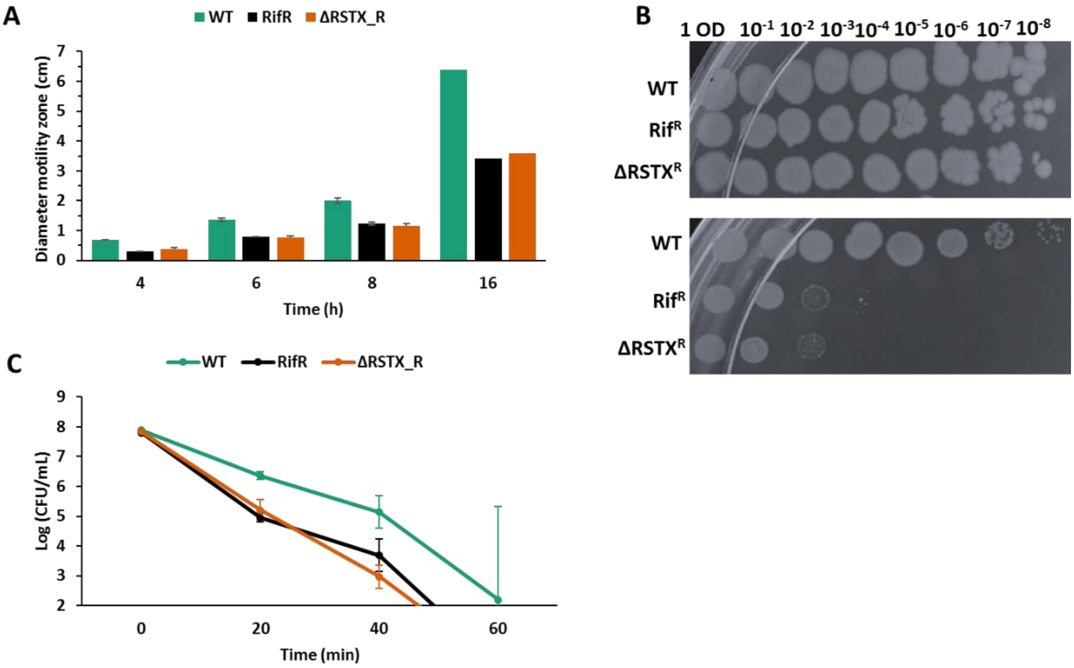


**Figure S2. Phenotypic characterisation of *V. vulnificus* CMCP6 wild-type (green), Rif$^{R}$ parental strain (black) and the ΔRSTX mutant (orange) in LBN.** A) Motility assay on tryptone motility plates. The strains were stabbed on motility agar plates and incubated at 37$^{o}$C. The motility zone was measured at several time points (4 h, 6 h, 8 h and 16 h). Three biological triplicates for each strain were tested (except at 16 h) and the reported values are the mean of the three replicates. B) Growth assessment in hyperosmotic stress conditions. The strains were first diluted to OD$_{600}$ = 1 and then 10-fold serial dilutions were performed up to 10$^{-8}$ and each dilution was spotted on LBN (top panel) and LBN + 0.4 M NaCl (bottom panel) and incubated at 37$^{o}$C. C) Survival assay in LBN at pH 4. The strains were incubated at 37$^{o}$C and survival was assessed, through plate counting, at four different time points (0 min, 20 min, 40 min and 60 min). Y-axis starts at Log (CFU/mL) = 2, in proximity of the detection limit value (2.4).

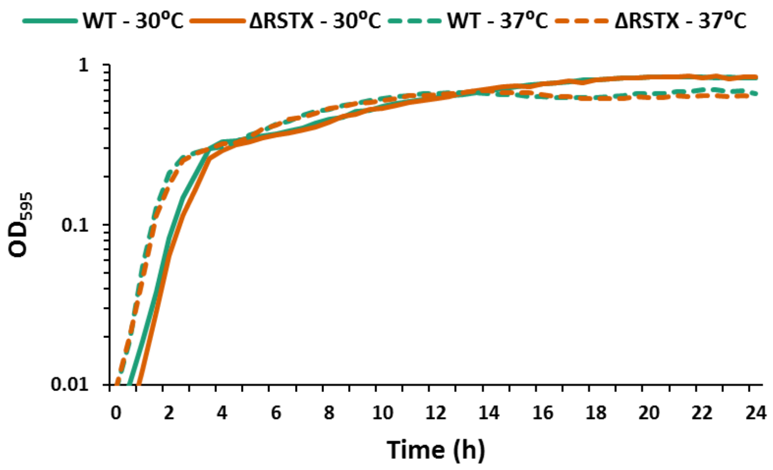


**Figure S3. Growth characterisation of *V. vulnificus* CMP6 wild-type (green) and the ΔRSTX mutant (orange) in LBN.** Growth curve in LBN, at 30$^{o}$C (continuous line) and at 37$^{o}$C (dashed line). $OD_{595}$ was measured every 30 min for 24 h. Each curve is the mean of three biological replicates. Y-axis starts at the OD value of the inoculum (0.01).
